# Encoding and context-dependent control of reward consumption within the central nucleus of the amygdala

**DOI:** 10.1101/2023.06.28.546936

**Authors:** Kurt M. Fraser, Tabitha H. Kim, Matilde Castro, Céline Drieu, Yasmin Padovan-Hernandez, Bridget Chen, Fiona Pat, David J. Ottenheimer, Patricia H. Janak

## Abstract

The ability to evaluate and select a preferred option among a variety of available offers is an essential aspect of goal-directed behavior. Dysregulation of this valuation process is characteristic of alcohol use disorder, with the central amygdala being implicated in persistent alcohol pursuit. However, the mechanism by which the central amygdala encodes and promotes the motivation to seek and consume alcohol remains unclear. We recorded single-unit activity in male Long-Evans rats as they consumed 10% ethanol or 14.2% sucrose. We observed significant activity at the time of approach to alcohol or sucrose, as well as lick-entrained activity during the ongoing consumption of both alcohol and sucrose. We then evaluated the ability of central amygdala optogenetic manipulation time-locked to consumption to alter ongoing intake of alcohol or sucrose, a preferred non-drug reward. In closed two-choice scenarios where rats could drink only sucrose, alcohol, or quinine-adulterated alcohol with or without central amygdala stimulation, rats drank more of stimulation-paired options. Microstructural analysis of licking patterns suggests these effects were mediated by changes in motivation, not palatability. Given a choice among different options, central amygdala stimulation enhanced consumption if the stimulation was associated with the preferred reward while closed-loop inhibition only decreased consumption if the options were equally valued. However, optogenetic stimulation during consumption of the less-preferred option, alcohol, was unable to enhance overall alcohol intake while sucrose was available. Collectively, these findings indicate that the central amygdala processes the motivational value of available offers to promote pursuit of the most preferred available option.

## INTRODUCTION

Selecting the best available option requires an assessment of the rewards currently available and a comparison of their relative values to appropriately direct motivated behavior. In substance use disorders, motivation becomes focused onto the pursuit of drug rewards despite the availability of other, perhaps more optimal, options^1–3^. The central amygdala (CeA) has been implicated in these processes as lesions or inactivation of this nucleus reduce drug self-administration and optogenetic stimulation of the CeA can promote choice of a stimulation-paired option over an otherwise equivalent reward^4–14^. However, these investigations have primarily been conducted with only one outcome type available, be it drug or natural rewards, and preclude an assessment of the contributions of the CeA to the dynamic process of choosing which outcome to pursue^15,16^.

The CeA is a candidate for outcome valuation, including ingested outcomes, as it receives privileged input from taste processing brainstem, thalamic, and cortical structures^17–20^. Previous work identified robust responses of CeA neurons to ingested outcomes^21–26^ and found pharmacological and optogenetic stimulation of some CeA cell populations increases intake of food or liquids^21,27,28^ further supporting a role for the CeA in a valuation process. While this work has begun to elucidate the means by which CeA neurons contribute to reward consumption more generally^21,22^, less is known about how neural activity within the CeA is related to alcohol consumption and choice *in vivo*. Oral alcohol shares sensory properties with food, but also has pharmacological properties that themselves profoundly impact CeA circuitry. Acute and, especially, chronic alcohol exposure alters CeA gene expression and synaptic physiology^29–33^. In addition, learning-dependent changes as a result of associating the taste of alcohol with its pharmacological effects may impact CeA function^32,34^. Of note, the taste of alcohol selectively activates mTORC1 signaling in the CeA and prelimbic and orbitofrontal cortices of alcohol-experienced rats and this activation of the CeA is necessary for the taste of alcohol to motivate alcohol-seeking^34^. Recently, Torruella-Suárez and colleagues demonstrated that manipulation of CeA neurotensin neurons in mice alters both alcohol and sucrose intake, further implicating this area in the regulation of alcohol intake^11^.

Here we examined the contributions of CeA neuronal activity to alcohol consumption in rats with a prior history of intake of alcohol at high levels to better understand how alcohol consumption may engage this region. In addition, we tested the contributions of CeA activity to reward choice, using optogenetic manipulation of the CeA time-locked to consumption to alter the choice between pursuit of alcohol over other options. We reveal robust neural responses to alcohol and natural reward consumption in the CeA and find that optogenetic increases or decreases of CeA neural activity can increase or decrease reward consumption, respectively. Of note, while the motivation to consume an outcome is modulated by CeA neural activity, the impact of CeA activation or inhibition is constrained by subjects’ preference between reward and drug options currently available in the environment. Collectively our findings suggest that the CeA acts as a motivational filter to focus reward and alcohol pursuit in a context-dependent manner.

## RESULTS

### Encoding of alcoohl consumption within the central amygdala

To evaluate potential correlates of alcohol consumption we pre-exposed rats in their homecage to 15% alcohol for 24 hours a day, 3 days a week, for 4-5 weeks^35,36^. This allowed rats to become familiar with the taste of alcohol and to associate alcohol consumption with its pharmacological effects, and, as well, to develop a propensity to consume relatively high levels (**Figure 1A**). Following initial homecage exposure, rats were implanted with drivable bundles of electrodes targeted to the CeA (**Figure 1B-C**) and neural activity was measured in freely-moving rats during alcohol consumption from a reward port situated in a standard behavioral chamber. Rats could enter the port to drink alcohol from a small receptacle that was primed with alcohol at session start. Once rats entered the port, a cumulative presence of 2 seconds activated the pump to deliver 0.1 mL of alcohol, a volume chosen to approximate the volume of alcohol typically consumed within 2 seconds (**Figure 1D**). Thus, we approximated continuous liquid delivery with no breaks when subjects chose to drink. We recorded 208 well-isolated single-units in 5 rats from the CeA during the consumption of alcohol with an average firing rate of 3.34 ± 0.198 (mean + s.e.m) Hz. During these sessions rats made on average 29.54 ± 2.76 port entries, 332.3 ± 43.78 licks, and drank on average 0.75 ± 0.07 g/ kg of alcohol. This level of alcohol consumption in this 30-40 minute timeframe produces blood alcohol contents of approximately 17.37 ± 4.24 mg/dL (data from a cohort of rats solely tested for alcohol consumption; n=17). We identified diverse responses to consumption-relevant behaviors (**Figure 1E**), primarily decreases in firing at approach (**Figure 1 F-H**) to the alcohol and exit (**Figure 1 M-O**) from the alcohol-containing port. Close inspection of the responses of single neurons that were significantly modulated by the first lick suggested that these neurons may be rhythmically modulated by the protrusion and retraction of the tongue, leading us to conduct further analysis to characterize CeA neural correlates of alcohol consumption.

**FIGURE 1:**
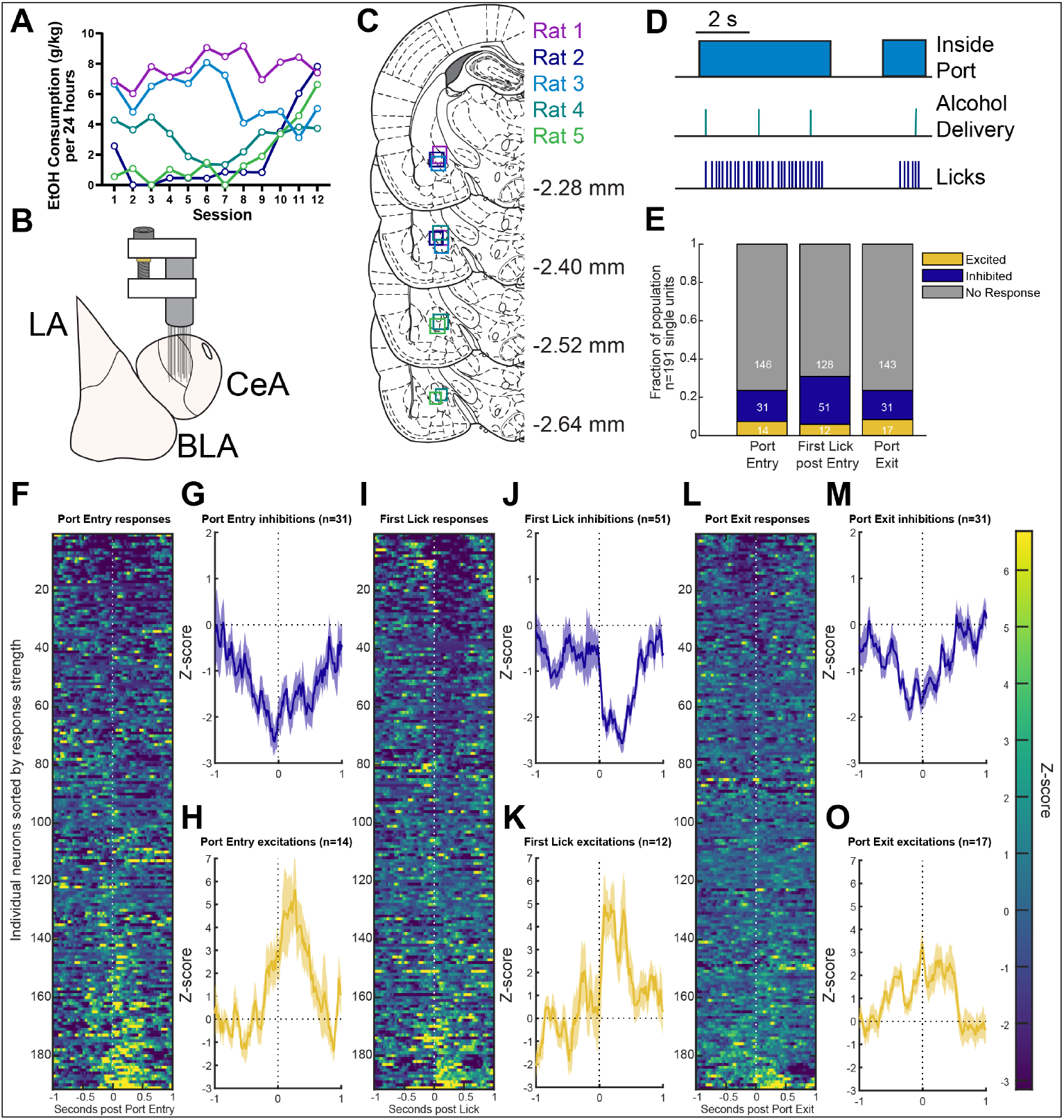
Identifying correlates of alcohol consumption in the central nucleus of the amygdala. **A)** Consumption of alcohol during a 4-week every other day, 24-hour availability to 15% alcohol on an intermittent access schedule expressed in g alcohol consumed per kg body weight. **B)** Schematic of recording strategy of 16 50-μm diameter tungsten wires in a drive targeted to the central amygdala. **C)** Recreation of recording sites from each of the five rats. **D)** Overview of task design. Sessions started with a prime of alcohol delivery in the port, and every cumulative 2 seconds spent in the port thereafter triggered a new delivery of alcohol. **E)** Proportion of neurons significantly excited or inhibited by task-relevant events. **F)** Heatmap of z-scored responses for each neuron recorded sorted by the strength of excitation to port entry. **G)** Average z-scored response of all neurons that were identified as being significantly inhibited around port entry. **H)** Average z-scored response of all neurons that were identified as significantly excited around port entry. **I-K)** Same as **F-H** but for the first lick post port entry. **L-O)** Same as **F-H** but for port exit. Heatmaps are sorted individually for each event. Traces indicate mean z-scored response with overlaid bands indicating ± 1 standard error of the mean.

To better relate CeA activity to the ongoing consumption of alcohol, we first analyzed the average lick rate for each recording session. Rat licking is highly stereotyped and typically averages 7 licks per second but can vary within the 5-10 Hz frequency range^37,38^. We observed similar average lick rates within this frequency band during alcohol consumption, averaging around 7 Hz across sessions and between rats (**Figure 2A**). Next, we assessed whether the firing of each of the 208 CeA neurons was modulated by rhythmic licking. To do so, we computed the firing phases of each spike relative to the lick cycle (phase 0 represents the contact of the tongue with the fluid delivery port; **Figure 2B**) and tested whether these firing phases were uniformly distributed across the lick cycle. Neurons with p-values below 0.01 were considered lick-modulated (Rayleigh test for non-uniformity; **Figure 2C**). We observed significant lick-modulation in the CeA as rats consumed ethanol with 24% of the population exhibiting significant lick-modulation (**Figure 2D**). At the population level, the large majority of lick-modulated neurons maximally fired early in the lick cycle, right after contact with the alcohol during the retraction of the tongue (**Figure 2E-F**). Consistent with a previous description of lick coherent firing of amygdala neurons^39^, the distribution of the preferred firing phases showed that ∼84% (32/38) of the lick-modulated neurons preferentially fired between 0 and π, with an overall mean direction of π/2 (**Figure 2G**). This suggests that, in addition to encoding the approach to alcohol, CeA neurons are phase locked to the lick rhythm during alcohol consumption and preferentially fired during the retraction of the tongue into the mouth right after the retrieval of alcohol from the environment.

**FIGURE 2:**
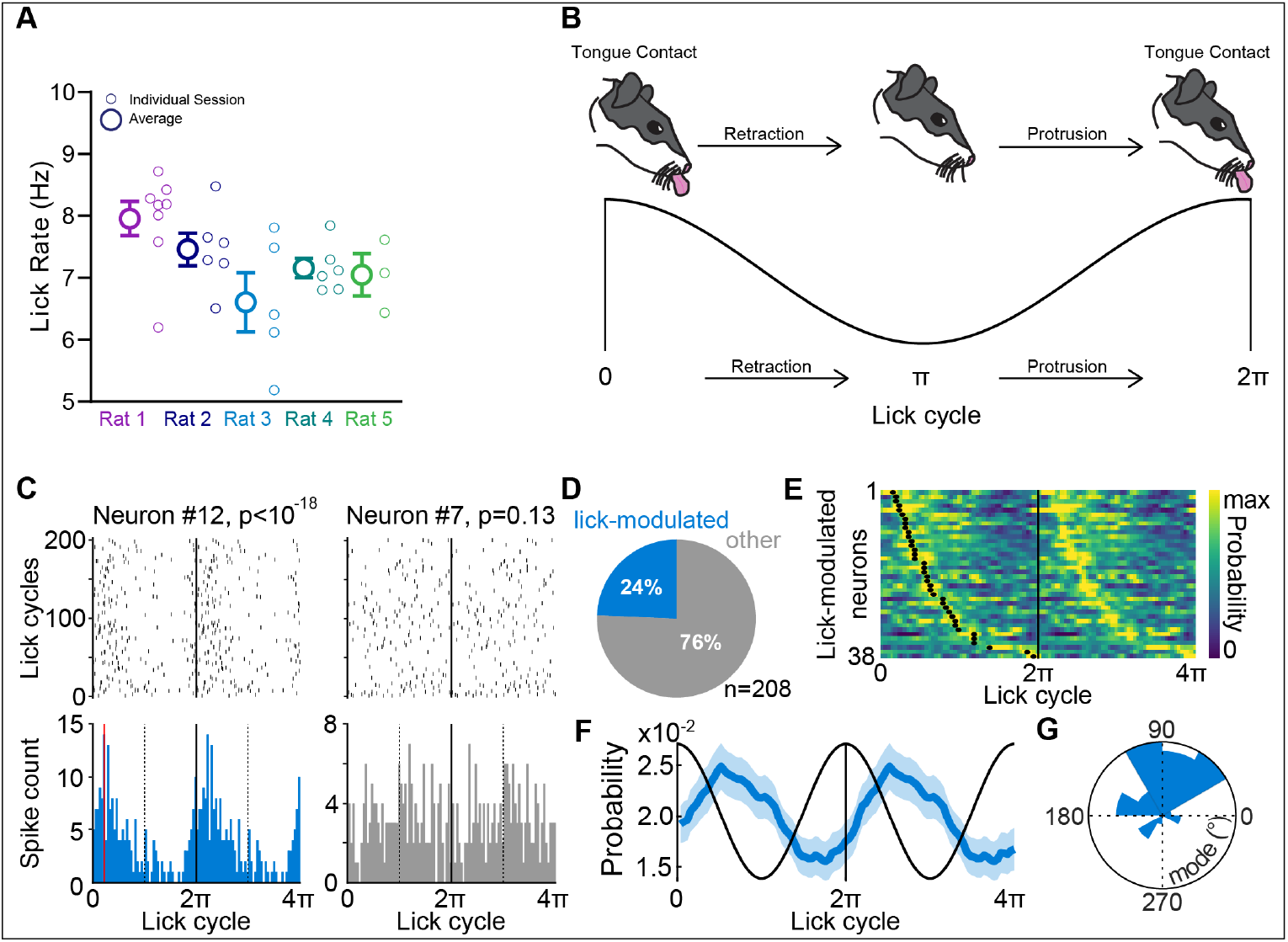
Central amygdala neurons are modulated by licks during the consumption of alcohol. **A)** Average lick rates during the consumption of 10% alcohol for each rat during recording session. Smaller symbols indicate lick rate for each individual session. **B)** Schematic representation of an individual lick cycle and its analogous sinusoidal rhythm. Times 0 and 2π represent contact with the liquid reward. **C)** Spike rasters (top) and histograms (bottom) during lick cycles for two example neurons recorded in the same session. A lick cycle is defined as the time between two consecutive contacts with the fluid delivery port (see Methods). The p-value of Rayleigh test is indicated. **D).** Proportion of neurons significantly modulated by licks (Rayleigh’s test with p-value < 0.01). **E)** Heat map of spike probability during lick cycles of lick-modulated neurons. Black dots indicate the preferred firing phases (i.e. modes). **F)** Average spike probability of lick-modulated neurons across lick cycles (mean±s.e.m.). **G)** Circular histogram of the preferred firing phases (V test against 90°, n=38 lick-modulated neurons, V_38_ =19.92, p<10^−5^).

We also recorded neurons in the CeA during the consumption of a concentration of sucrose isocaloric to 10% ethanol. We observed similar overall patterns of activity at the time of sucrose approach (**Supplement 1**) and lick-modulation in the CeA during sucrose consumption (**Supplement 2**) suggesting common codes for consumption of rewards in the CeA. Interestingly, we observed more lick-entrainment in the CeA of rats consuming sucrose than for the consumption of ethanol (**Supplemental Figure 2G**). However, there was no difference in the preferred phase of the lick cycle for lick-modulated CeA neurons during alcohol and sucrose consumption (**Supplemental Figure 2H**). This observation suggests that the degree of lick-modulation in the CeA neuronal population is influenced by reward value. Together, these data indicate that the CeA is modulated by the pursuit of reward and is strongly engaged during the ongoing consumption of both natural and drug outcomes.

### Optogenetic excitation of the central amygdala biases reward and alcohol choice but is filtered by preference

We observed correlates of consumption and motivation to pursue alcohol in the CeA and sought to understand the functional consequences of this activity. To do this, we expressed either the excitatory, blue-light activated opsin ChR2 or a control fluorescent protein GFP in the CeA of rats (**Figure 3A-B**). Rats became familiar with the taste and pharmacological properties of alcohol in their homecage by allowing rats 24 hours of access to 15% alcohol 3 days a week for 5 weeks (**Figure 3C**). We then gave rats 30-minute access to offers of pairs of solutions, comprised of different combinations of 10% alcohol, isocaloric 14.2% sucrose, and 10% alcohol adulterated with 100 μM quinine, an outcome commonly used to test for compulsive consumption of alcohol^40–42^, in a modified homecage. For each test session, licks were recorded from each of the two bottles. Licks made on one of the bottles triggered delivery of a 1 s, 20 Hz train of 8-12 mW blue light bilaterally into the CeA (**Figure 3D**).

**FIGURE 3:**
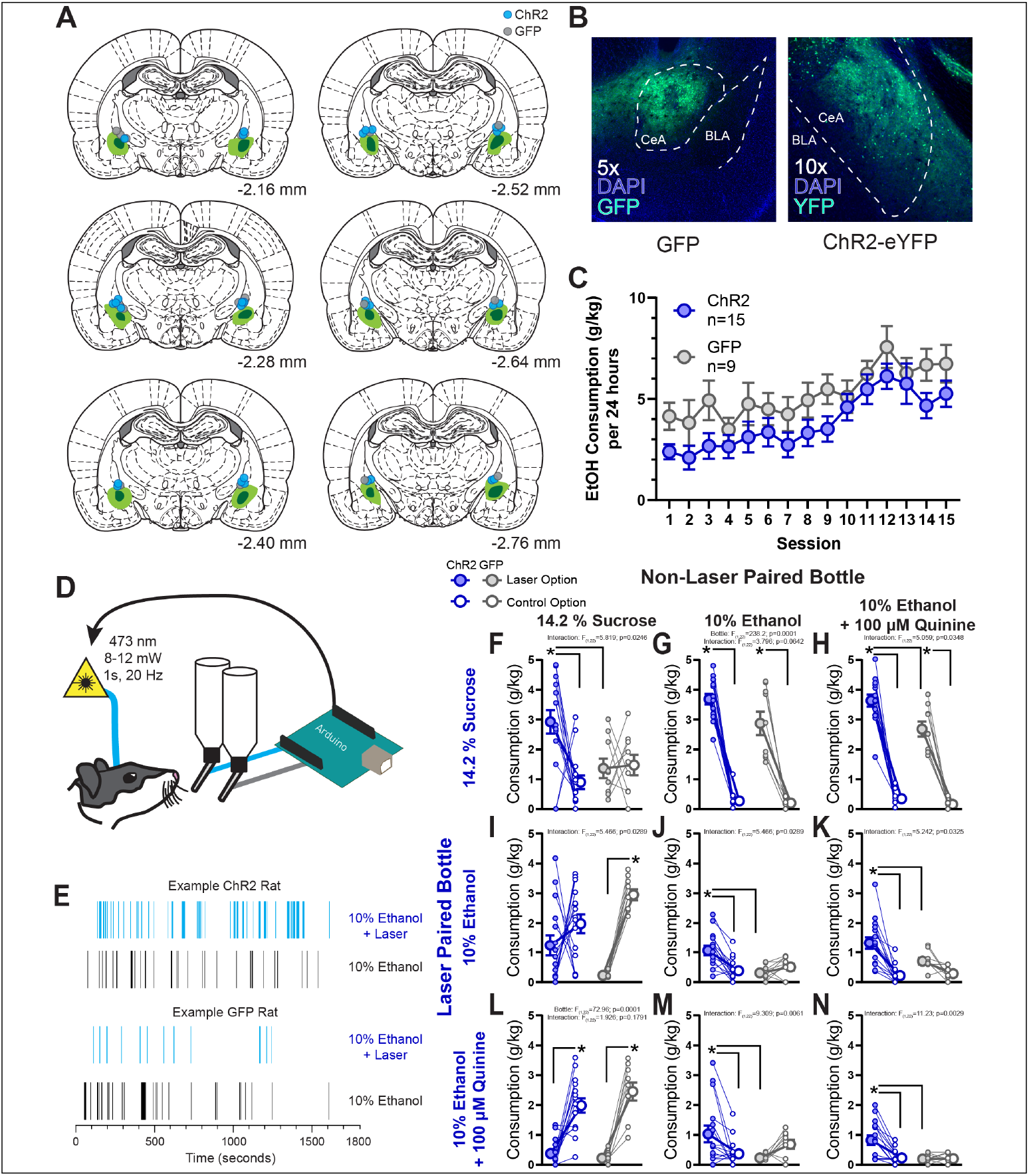
Context-dependent enhancement of reward consumption resulting from closed-loop optogenetic stimulation of the central amyg-dala. **A)** Reconstruction of the maximal (light green) and minimal (dark green) viral expression of either hsyn-ChR2-eYFP or hsyn-GFP within the central amygdala. Blue dots indicate fiber tips for rats with ChR2 and grey dots indicate fiber tips for rats with GFP. **B)** Example images of expression of GFP and ChR2-eYFP within the central amygdala (green) and nuclear staining with DAPI in blue. **C)** Homecage alcohol consumption (in g alcohol per kg body weight) on an every-other day, 24 hour intermittent access schedule to 15% alcohol. Consumption in the homecage did not differ between rats with ChR2 or GFP expression in the central amygdala (F_1,22_=2.658, p=0.1172). **D)** Rats were allowed to freely direct their consumption between two bottles containing a variety of liquid rewards. Licks were recorded on each bottle by an Arduino and in turn the first lick each second on one of the bottles would result in a 1 s, 20 Hz train of blue light delivered bilaterally to the central amygdala. **E)** Example lick rasters from a session in which both bottles contained 10% alcohol for a representative ChR2 rat and a representative GFP rat. Consumption of each solution in g consumed per kg body weight. Graphs are organized with the most valued option at the top and leftmost position and the least valued option at the bottom and rightmost position. Comparisons between bottles containing the same offer tile the diagonal, bottles above diagonal are tests in which the more valued option was blue-light paired and tests below the diagonal are when blue-light was paired with the less valued option. Blue symbols represent ChR2 rats, grey symbols indicate GFP rats. Filled symbols indicate the bottle that resulted in blue light delivery, open symbols the other bottle that did not trigger any light delivery. Large symbols indicate group means ± 1 standard error of the mean and small symbols represent individual rats. * p<0.05 for post hoc comparisons made only when a significant main effect of bottle or an interaction between virus group and bottle were observed.

We found that rats expressing ChR2 in the CeA would consume more of the laser-paired option if the other non-laser paired option was identical (**Figure 3E** for example licking behavior at test). This was true for sucrose (**Figure 3F**), alcohol (**Figure 3J**), and quinine-adulterated alcohol (**Figure 3N**). We examined the microstructure of consumption to determine the psychological mechanisms underlying this effect, focusing on clusters of licks (drinking bouts) and the number of licks within a cluster. Cluster number is typically taken to reflect motivation for a particular substance while lick number per cluster is correlated with palatability^37,43–45^. In GFP control rats, mean cluster number was correlated with palatability with sucrose>alcohol>quinine-adulterated alcohol, as expected. We observed that stimulation primarily increased the number of drinking bouts made to the laser-paired option, but rarely increased the average number of licks within each cluster, especially in these tests where the options were otherwise identical (**Supplement 3**). This pattern of increased clusters, or number of drinking bouts, suggests that CeA stimulation enhanced the motivation to pursue and consume the laser-paired option but did not enhance the perceived palatability of the option^37,43,45^.

Next, we asked whether CeA stimulation would alter the choice rats made between two different options. Interestingly, we found that if the laser-paired option was preferred by the rats, then stimulation enhanced consumption of the preferred reward even further above that observed in control rats. For example, optogenetic stimulation enhanced intake of sucrose and of alcohol when each of these was paired with quinine-adulterated alcohol (sucrose over alcohol-quinine **Figure 3H**; alcohol over alcohol-quinine **Figure 3K**). While the interaction between bottle and group did not reach significance for laser-paired sucrose consumption with alcohol as the other option (**Figure 3G**), this is potentially due to a ceiling effect of consumption in this time-limited test. Nonetheless, stimulation paired with reward ingestion in all of these cases significantly increased motivation to consume the laser-paired option as evidenced by increased drinking bouts, but did not appear to alter the hedonic value of the laser option (**Supplement 3**).

In contrast, when CeA stimulation was paired with the non-preferred option, and sucrose was the other option available in the choice, stimulation did not reliably increase consumption of the laser-paired option. This was especially apparent in the case of alcohol-quinine vs sucrose, where stimulation had no impact on intake (**Figure 3L**). Interestingly, stimulation also failed to increase alcohol intake over sucrose when analyzing intake (**Figure 3I**), but stimulation did increase the mean number of lick clusters for alcohol vs sucrose (**Supplement 3N**), suggesting an ability of CeA stimulation paired with alcohol consumption to promote choice of that option over other possible rewards. Elevated motivation to consume alcohol in this test for ChR2 rats was not related to overall measures of propensity to drink alcohol, as the correlation between alcohol intake on the laser-paired bottle and homecage alcohol consumption was not significant (r=-0.4534, p=0.0896).

Finally, when alcohol was the control offer and quinine-adulterated alcohol was stimulation-paired, ChR2 rats drank more quinine-adulterated alcohol (**Figure 3M**), and increased the number of lick clusters (**Supplement 3S**), perhaps representative of reduced sensitivity to punishment in some individuals^40^. Of note, there was no correlation between alcohol-quinine intake when it was laser-paired and alcohol was available and the final homecage alcohol drinking session (r=-0.3988, p=0.1409) for ChR2 rats.

This pattern of findings suggests that the CeA compares the value among available options and directs motivation to the most preferred offer, but that underlying preferences that inform directed motivation may be independent of the CeA. This conclusion is based on the above choices with alcohol, a frequently less-preferred outcome in non-dependent rats, and may not generalize to comparisons among non-drug outcomes. To better isolate the possibility of selective enhancement of motivation to preferred outcomes, we offered rats the choice between sucrose and maltodextrin, two isocaloric sweet rewards that are equally consumed when presented in isolation, but differ in that rats prefer to consume sucrose when given a choice^38,46–48^. When both bottles contained maltodextrin, ChR2 rats consumed more of the maltodextrin leading to CeA stimulation (**Supplement 4**). When CeA stimulation was paired with sucrose consumption when maltodextrin was available, CeA stimulation enhanced sucrose consumption above that observed in control rats as we observed with choice between laser-paired sucrose and alcohol. Surprisingly, when CeA stimulation was paired with maltodextrin consumption and the more preferred sucrose was the other option, ChR2 rats enhanced maltodextrin consumption, reversing preference between the two rewards (**Supplement 4**). These findings collectively indicate that enhancing CeA neural activity can strongly bias reward choice resulting in increased consumption of the laser paired option, even for drugs of abuse, especially if that option is of greater or similar value than other offers.

### Optogenetic stimulation of the central amydala is reinforcing

CeA stimulation selectively enhanced consumption of the most valuable available reward, highlighting a possibility for an innate reinforcing property of CeA stimulation that could promote responding absent physical reward. Evidence for such a primary reinforcement signal in the CeA has been mixed^9–14,49,50^. We first gave rats the option to drink water that was paired with CeA stimulation. As before, stimulation increased the consumption of laser-paired water, which was surprising given rats were not water-restricted (**Figure 4A-C**; response: F_1,22_=12.56, p=0.0018; group: F_1,22_=10.24, p=0.0041; interaction: F_1,22_=12.98, p=0.0016; stimulations: t_14_=4.491, p=0.0005). We then asked if rats would respond without reward available to earn CeA stimulation by offering empty bottles. We observed rats would lick on the empty bottle to earn CeA activation (**Figure 4E-G**; response: F_1,22_=30.45, p < 0.0001; group: F_1,22_=30.59, p < .0001; interaction: F_1,22_=32.05, p < 0.0001; stimulations: t_14_=6.377, p<0.0001). We then wanted to rule out potential extinction effects contributing to this behavior as the rats had extensive experience in drinking from bottles in the testing apparatus. We placed rats into a novel operant behavioral chamber with two nosepoke ports available. Responses in one nosepoke led to CeA stimulation. Rats acquired this novel response and worked to earn CeA stimulation (**Figure 4I-K**; response: F_1,22_=8.019, p = 0.0097; group: F_1,22_=7.92, p=0.0101; interaction: F_1,22_=7.807, p=0.0106; stimulations: t_14_=4.674, p=0.0004). A subset of rats (n=7 ChR2, n=3 GFP) were tested for intracranial self-stimulation prior to homecage ethanol drinking to assess whether the consistent self-stimulation we observed could result from ethanol-induced plasticity. However, even alcohol-naive rats exhibited self-stimulation of the CeA (**Figure 4D, 4H, 4L**; water: t_6_=3.23, p=0.0178; empty:t_6_=2.959, p=0.0249; nosepoke: t_6_=3.809, p=0.0088). These data indicate there is a reinforcing property of CeA stimulation itself, but, in the case of external reward availability, this reinforcing property is filtered by preference among the available rewards.

**FIGURE 4:**
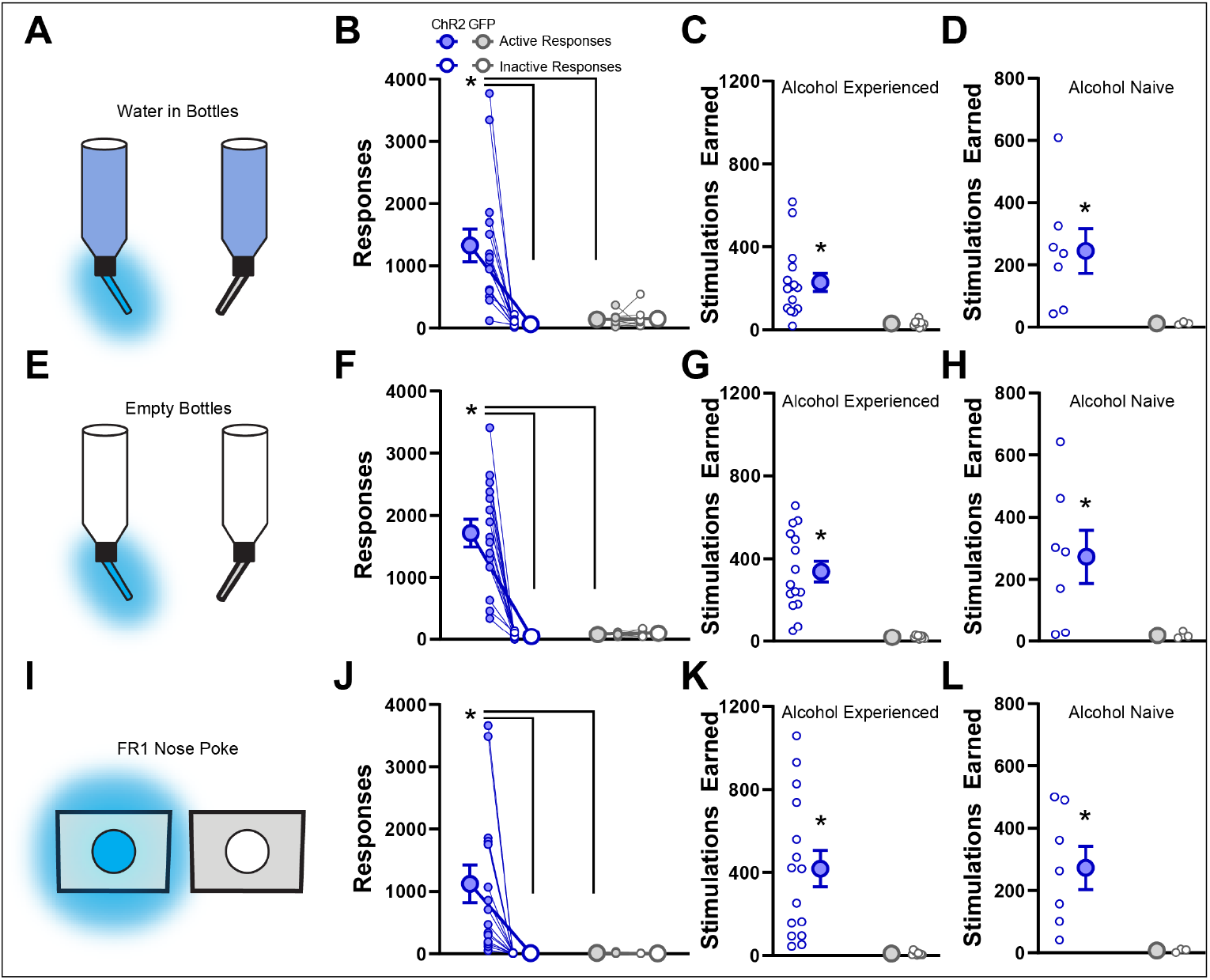
Optogenetic self-stimulation of the central amygdala is reinforcing regardless of alcohol experience. **A)** Rats were allowed 30 minutes of access to water-containing bottles in the modified homecage and the first lick to one bottle resulted in a 1s, 20 Hz train of blue light delivered bilaterally to the central amygdala. **B)** Rats with ChR2 in the central amygdala made significantly more licks to the stimulation-paired bottle than the control bottle and also more than GFP rats. **C)** Total stimulations earned during the session. D) Total stimulations earned for a subset of rats tested prior to any alcohol drinking experience. **E-H)** same as **A-D** but for a test in which both bottles were empty. ChR2 rats made significantly more licks to the stimulation paired bottle and earned significantly more stimulations than GFP rats. **I-L)** Same as **A-D** but instead of bottles, rats were placed in operant conditioning chambers and allowed to nose poke freely where the first poke each second in port resulted in a 1s, 20 Hz train of blue light delivered bilaterally into the central amygdala. Filled symbols indicate the bottle that resulted in blue light delivery, open symbols the other bottle that did not trigger any light delivery. Large symbols indicate group means ± 1 standard error of the mean and small symbols represent individual rats. * p<0.05 for post hoc comparisons made only when a significant main effect of bottle or an interaction between virus and bottle were observed.

### Optogenetic inhibition of the central amygdala reduces reward valuation

We observed that stimulation of the CeA can increase the motivation to pursue and consume rewards. We were then interested in examining the effects of inhibition of the CeA. To do this, we expressed either the inhibitory, green- and yellow-light activated opsin halorhodopsin (eNpHR) or a control fluorescent protein GFP in the CeA of rats (**Figure 5A-B**), and trained rats as above to choose from bottle pairs presented in 30 minute tests; for each test, licks were recorded from each of the two bottles and licks made on one of the bottles were associated with the delivery of a constant train of 15-20 mW green light bilaterally into the CeA until no lick was detected for 1 s (**Figure 5D**).

**FIGURE 5:**
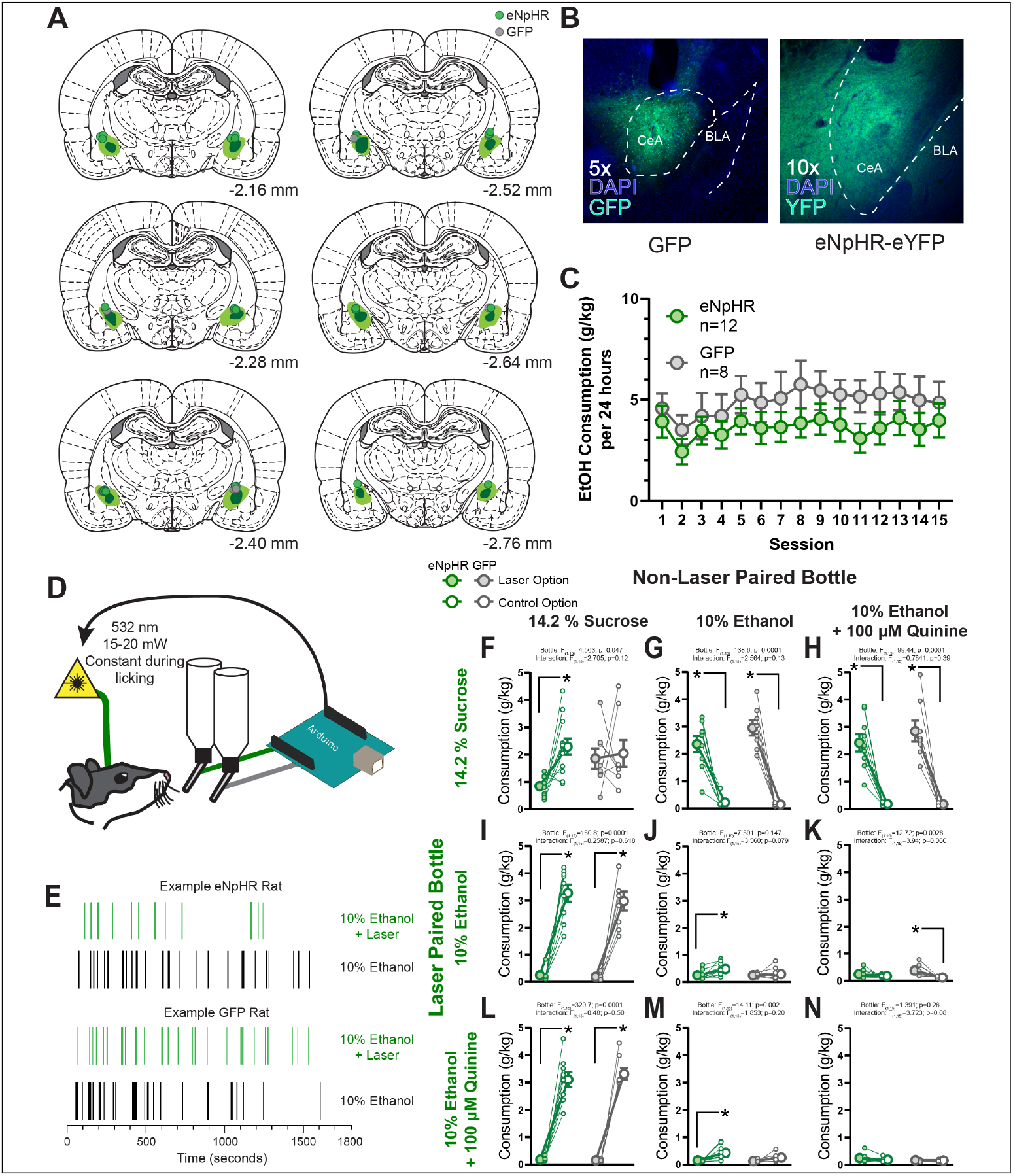
Context-dependent suppression of reward consumption resulting from closed-loop optogenetic inhibition of the central amygdala. **A)** Reconstruction of the maximal (light green) and minimal (dark green) viral expression of either hsyn-eNpHR-eYFP or hsyn-GFP within the central amygdala. Green dots indicate fiber tips for rats with eNpHR and grey dots indicate fiber tips for rats with GFP. **B)** Example images of expression of GFP and eNpHR-eYFP within the central amygdala (green) and nuclear staining with DAPI in blue. **C) H**omecage alcohol consumption (in g alcohol per kg body weight) on an every-other day, 24 hour intermittent access schedule to 15% alcohol. Consumption in the homecage did not differ between rats with eNpHR or GFP expression in the central amygdala (F_1,18_=1.695, p=0.2093). **D)** Rats were allowed to freely direct their consumption between two bottles containing a variety of liquid rewards. Licks were recorded on each bottle by an Arduino and in turn the first lick on one of the bottles would result in a constant train of green light delivered bilaterally to the central amygdala. **E)** Example lick rasters from a session in which both bottles contained 10% alcohol for a representative eNpHR rat and a representative GFP rat. symbols. **F-N)** Consumption of each solution in g consumed per kg body weight. Graphs are organized with the most valued option at the top and leftmost position and the least valued option at the bottom and rightmost position. Comparisons between bottles containing the same offer tile the diagonal, bottles above diagonal are tests in which the more valued option was green-light paired and tests below the diagonal are when green-light was paired with the less valued option. Green symbols represent eNpHR rats, grey symbols indicate GFP rats. Filled symbols indicate the bottle that resulted in green light delivery, open symbols the other bottle that did not trigger any light delivery. Large symbols indicate group means ± 1 standard error of the mean and small symbols represent individual rats. * p<0.05 for post hoc comparisons made only when a significant main effect of bottle or an interaction between virus group and bottle were observed.

Overall, the effects of optogenetic inhibition of the CeA on reward consumption were much less robust than of excitation. When examining choices among sucrose, alcohol, and alcohol-quinine, almost none of the two-way ANOVAs comparing virus group and bottle revealed significant interactions. We considered that reductions in already relatively low levels of alcohol intake would be difficult to detect due to floor effects, and therefore chose to conduct post hoc comparisons within each virus group across the two bottles to evaluate our a priori hypotheses based on our strong findings with optogenetic stimulation.

We found that rats with eNpHR in the CeA consumed less of the laser-paired option if the other non-laser paired option was identical (**Figure 5E** for example licking behavior at test). This was true for sucrose (**Figure 5F**) and alcohol (**Figure 5J**). Baseline consumption of quinine-adulterated alcohol was already low so there was no effect of inhibition (**Figure 5N**). We examined the microstructure of consumption to determine the psychological mechanisms underlying this effect. In cases where a significant decrease in intake was observed, we did not find a significant decrease in the number of clusters of licks made to the laser paired option but did find a decrease in the average number of licks within each cluster for sucrose, but not alcohol (**Supplement 5**). However, given that consumption was low, this decrease may be a result of floor effects of overall consumption from the limited 30-minute sessions.

We then asked whether CeA inhibition would alter the choice rats made between two different options. We found that if the laser-paired option was preferred by the rats, (e.g. sucrose over alcohol, alcohol versus quinine-adulterated alcohol) inhibition could not reduce consumption of the preferred reward below that observed in control rats (sucrose over alcohol **Figure 5G**; sucrose over alcohol-quinine **Figure 5H**; alcohol over alcohol-quinine **Figure 5K**). In addition, when CeA inhibition was paired with the non-preferred option, the laser had no effect on consumption (alcohol vs sucrose **Figure 5I**; alcohol-quinine vs sucrose **Figure 5L**; alcohol-quinine vs alcohol **Figure 5M**).

Once again, we offered rats the choice between sucrose and maltodextrin to compare the ability of optogenetic inhibition of the CeA to alter choice between two similarly-valued rewards. When both bottles contained maltodextrin, there was a borderline interaction between bottle and group, accounted for by lower consumption by eNpHR rats of maltodextrin paired with CeA inhibition (**Supplement 6B**). When CeA inhibition was paired with sucrose consumption when maltodextrin was available, CeA inhibition did not reduce sucrose consumption (**Supplement 6A**). When CeA inhibition was paired with maltodextrin consumption and the more preferred sucrose was the other option, both eNpHR and GFP rats consumed more sucrose (**Supplement 6C**), and consumption of maltodextrin was not significantly decreased below control levels. Unlike stimulation of the CeA, inhibition of the CeA only reduced reward preference in closed-choice scenarios. Taken together, CeA inhibition reduces consumption of the laser-paired option when the other option is of equal value but does not reverse preference between disparately valued rewards.

## DISCUSSION

Here we describe encoding of alcohol consumption in the central amygdala and provide a demonstration of conditions in which central amygdala stimulation and inhibition can alter alcohol and other reward preference. We report a phasic signal in central amygdala neurons during licking for both drug and natural rewards that is entrained to the lick cycle. In addition, when optogenetic manipulation of the central amygdala is time-locked to the consumption of rewards we reveal a context-dependent ability of stimulation and inhibition to alter consumption. In a context where alcohol is one of the available choices, only when the most preferred currently available reward is paired with stimulation does the central amygdala contribute to prolonged pursuit and consumption of either natural or alcohol outcomes. Further, optogenetic inhibition of the central amygdala only decreased consumption when options in the environment were the same (both bottles containing sucrose or alcohol). Together, these findings provide evidence that the central amygdala is a critical node in decision-making circuitry that integrates value-related information about available rewards to filter and refine motivation.

The CeA is historically known as critical for the scaling and expression of fear-related responses^51^, with the interaction between the CeA and primary taste centers in the brain receiving relatively less attention. The taste-related afferents into the CeA may allow for the integration of the hedonic properties of reward with relevant state- and history-dependent representations within the CeA. Indeed, afferents from the parabrachial nucleus and insular cortex to the CeA are essential for the avoidance of a previously appetitive tastant made aversive through pairing with gastrointestinal distress^52,53^. CeA inputs from the insula, either direct, or indirect via the basolateral amygdala, have been proposed to mediate assignment of taste value^27^. The CeA could then integrate this taste-information into relevant cell-type and projection-specific circuits to select the appropriate consummatory action and scale the degree of this response appropriately^21,22,52–58^.This integration of hedonic and motivational information in the CeA and its efferents to brainstem motor centers that control both the jaw and the initiation of movement towards a target of motivation situates the CeA as a limbic command center for consummatory behaviors^59,60^.

Our findings that rats will avidly work for optogenetic activation of the CeA further supports the role of CeA in positively-motivated behavior, and is in line with multiple reports in mice ^11,22,27,28,50^. That rats will self-stimulate the CeA suggests that the natural activation of at least some of these neurons can participate in a reinforcement process. However, our findings of alcohol- or sucrose-paired enhancement of intake cannot be ascribed to self-stimulation alone, since the amount of intake of solutions paired with stimulation was variable and depended on the specific options available.

For example, we found that CeA activation during alcohol consumption more than doubled alcohol intake when rats chose between two bottles, each containing some combination of alcohol and/ or alcohol with quinine. However, when an alcohol-containing bottle was paired with CeA activation and sucrose was available in the neighboring bottle, the effect was much weaker. Relative to control rats, mean intake of quinine-adulterated alcohol was not increased by optogenetic CeA stimulation when sucrose was also available, and, although the number of lick clusters for alcohol alone was significantly increased when sucrose was in the unstimulated bottle, total intake of alcohol was not significantly altered. Inspection of the values from individual rats clearly shows a large within-group variation on the impact of alcohol-paired CeA activation within the alcohol-sucrose choice. It could be that rats with a greater propensity to drink alcohol might be more easily shifted by optogenetic stimulation; yet we did not find a correlation with baseline drinking levels and drinking levels during optogenetic stimulation. However, other means to evaluate motivation for alcohol could reveal systematic behavioral underpinnings of the variation we observed. For example, some innate liability to aversion-resistant drinking that is not captured by overall levels of alcohol intake may contribute. It is possible then that individual variation and experience-dependent alterations in the coding of taste of rewards and their resultant value may dynamically influence the role of the CeA in choice. In addition, although no obvious patterns emerged, individual differences in virus expression within the extent of the CeA, or even within different cell types within the CeA, could also account for these distinctions, and should be better evaluated within alcohol choice drinking models.

The selective enhancement of consumption for the most preferred option is in contrast to a recent report that CeA stimulation can result in the choice of less-preferred cocaine over sucrose^12^. However, in that study, rats were required to sample each option at session start, unlike the current study. It is also important to note that sucrose requires oral consumption whereas cocaine is delivered intravenously, so the activation of CeA for a prolonged period during a cocaine infusion may be supraphysiological as the activity of the CeA during intravenous drug delivery is unclear. Our report of dynamic contributions of CeA stimulation time-locked to the consumption of rewards equated the cost, response effort, and consummatory behavior associated with each of the two outcomes in a given choice providing confidence that consumption altered by CeA stimulation was due to an acute change in the rats’ motivation.

In agreement with prior work^21,58^, our finding that stimulation of the CeA impacted consumption primarily by increasing the number of lick clusters made to the stimulation-paired bottle supports a role for CeA activity in motivation to consume the alcohol and sucrose outcomes. We cannot discount an influence of CeA neural activity on palatability, although the evidence to support this was weaker, limited to an effect of sucrose-paired CeA inhibition on the number of licks rats made in each cluster. If further work supports a role on outcome palatability, this might suggest that the CeA can separately modify the motivation to consume and the palatability of orally-ingested rewards.

The CeA is profoundly impacted by prior experience with drugs of abuse and in particular alcohol. The induction of physical dependence on alcohol in rodents, often via forced alcohol vapor exposure, recruits the CeA to play an essential role in escalating alcohol drinking, alcohol self-administration, and withdrawal-related anxiety-like behaviors^61,62^. These alcohol dependence-induced alterations in CeA circuitry have suggested a limited contribution of the CeA to alcohol-seeking and taking only after an individual has met a threshold of drug consumption or is made dependent^1,63^. In our experiments, rats volitionally drank alcohol in the homecage, which does not typically induce dependence, yet we found that stimulation of the CeA could promote the consumption of bitter quinine-adulterated alcohol despite the availability of a more preferred non-adulterated alcohol. This, along with the inability of CeA stimulation to flip preference for alcohol if sucrose was available, suggests a broader role for CeA circuits in evaluating currently available rewards and directing motivation to the most desirable option prior to the manifestation of physical dependence^11,64^. Moreover, despite low overall levels of alcohol consumption, inhibition of the CeA reduced consumption of inhibition-paired alcohol, similar to what has been reported for cocaine choice^13^. The ability of the options available to alter the effects of CeA activation and inhibition indicates that the CeA computes a relative comparison among options. This ultimately provides a potential mechanism for experience and dependence to impinge upon within the CeA to promote maladaptive and inappropriate alcohol-seeking and drinking. It will be important in the future to understand how alcohol-induced alterations of CeA might impact the control of alcohol intake we have demonstrated here.

The CeA is comprised of a number of diverse cell types, including neuropeptide-producing neurons, which we did not account for in these studies that likely contribute to the enhancement of motivation to seek alcohol and rewards. For instance, CeA neurotensin-expressing neurons that project to the parabrachial nucleus can control alcohol intake^11^, and prior studies using natural rewards have implicated prepronociceptin-, somatostatin-, and 5-HT2aR-positive CeA neurons in the promotion of food intake more generally^21,22^. Acute and chronic alcohol consumption alters CeA GABAergic and glutamatergic signaling, as well as expression of multiple neuropeptides such as CRF and NPY^14,29,30^. It will be critical, then, to understand how these diverse populations of cells in the CeA interact to promote alcohol-seeking in a dynamic environment where individuals have an array of desirable options available. Additionally, our studies only included male rats but it will be important in the future to dissect potential sex differences in the contribution of the CeA to alcohol choice^65^. Collectively, our findings suggest the CeA is a critical component of decision-making circuitry that interacts with motivation, preference, and experience to guide the pursuit and consumption of rewards.

## METHODS

### Animals

Male Long-Evans rats weighing 250-275 g and approximately 60 d of age upon arrival were obtained from ENVIGO (Frederick, MD; n=66) or were bred in our laboratory (n=5). Rats were single-housed in a temperature- and climate-controlled vivarium on a 12-h light:dark cycle. Rats were left undisturbed for at least one week in the vivarium before the beginning of behavioral training, alcohol exposure, or surgery. Water and food was available ad libitum and rats were provided with paper shredding enrichment in the homecage. Experimental procedures took place during the light phase of the light:dark cycle. All procedures were conducted in accordance with protocols approved by the Animal Care and Use Committee at Johns Hopkins University.

### Reward solutions

Ethanol was prepared fresh from 200 proof stock solution and diluted in tap water to either 15% by volume for homecage exposure or 10% by volume for electrophysiology and optogenetic experiments. Sucrose (Thermo Fisher Scientific) and maltodextrin (SolCarb, Solace Nutrition) were prepared as 14.2% solutions in tap water by weight. Quinine-adulterated ethanol was prepared by adding quinine salt to a solution of 10% ethanol to achieve a concentration of 100 μM.

### Surgical procedures

Rats were induced into a surgical plane of anesthesia by inhalation of 5% isoflurane and then maintained at 2-3% isoflurane for the duration of the surgical procedures. For rats in the electrophysiology experiments (n=12) a 1 mm craniotomy was made unilaterally above the central amygdala (AP: -2.4; ML: -4.2 relative to bregma) and 6-8 screws were placed in the skull for anchoring of the implant and one was selected as the screw for the ground wire. A custom-printed microdrive containing a bundle of 16 50 μm tungsten wires and 2 silver ground wires was then lowered slowly to the central amygdala (DV: -7.8 relative to bregma), the ground wires were wrapped around a skull screw, and the drive was secured to the skull with dental cement. For optogenetic experiments, rats received infusions of 500 nL of AAV5-hsyn-ChR2-eYFP (n= 22; Addgene 26973; 1.7 × 1013 viral particles per mL), AAV5-hsyn-eNpHR3.0-eYFP (n=12; Addgene 26972; 1.0 × 1013 viral particles per mL), or AAV5-hsyn-GFP (n=20; Addgene 50465; 1.2 × 1013 viral particles per mL) bilaterally into the central amygdala (AP: -2.4; ML: ±4.0; DV: -7.8 relative to bregma) at a rate of 100 nL/minute through a 31-gauge gastight Hamilton syringe attached to a Micro4 Ultra Microsyringe Pump 3 (World Precision Instruments) with a 10 minute waiting period prior to the removal of the needle. Rats then received with 300 μm diameter optic fiber bilateral implants aimed 0.3 mm above the site of virus infusion (DV: -7.5). Optic fiber implants were secured to the skull with dental cement and 4 skull screws. Rats received an injection of carprofen (5 mg/kg s.c.) immediately following surgery and were allowed to recover for at least 10 days.

### Histology

Rats were deeply anesthetized with sodium pentobarbitol. For rats with electrode implants, final electrode sites were marked by briefly passing a DC current through each electrode. All rats were then perfused with 4% paraformaldehyde and brains extracted and post-fixed for 24 hours at 4C. Brains were cryoprotected in 30% sucrose in 0.1M NaPB for 2-3 days, sliced on a freezing cryostat (Leica), and 50 μm sections were collected. Electrode locations were visualized by staining with cresyl violet. The locations of optical fiber tips and virus expression were visualized with immunohistochemistry. Briefly, slices were washed in 0.1M PBS and blocked in 10% normal donkey serum in 0.1M PBS for 30 minutes and then incubated at 4C overnight with primary antibody (mouse anti-GFP at 1:1500; Invitrogen A1120). The following day sections were washed in PBS and then incubated for 2 hours at RT in secondary antibody (Alexafluor 488 donkey anti-mouse at 1:200; Invitrogen A21202) following which they were washed, mounted onto slides, stained with DAPI (Vectashield; VWR H-150) and imaged on a fluorescence microscope (Zeiss).

### Homecage ethanol exposure

Rats were allowed to drink 15% ethanol freely in the homecage for 24 hours Monday, Wednesday, and Friday for either 4 weeks or 5 weeks depending on the experiment. Rats had free access to water the entire time via a Lixit spout in the homecage. Ethanol bottles were weighed before and after each drinking session and rat weights were recorded at the end of each drinking session.

### Ethanol and sucrose self-administration

For rats in the electrophysiological experiment, a modified self-administration protocol was used. A dish in a recessed port in a modified MedAssociates chamber was filled at the start of the session with 10% ethanol or 14.2% sucrose. During each 40-minute session, a 2 second cumulative presence in the reward-containing port resulted in the activation of a pump for 2 seconds. Based on pilot experiments we determined this matched the rate at which rats consumed the reward and resulted in the fluid dish almost always containing reward (∼0.1 mL per delivery). Licks were recorded from the reward-containing fluid dish via a custom-made lickometer, and port entries and exits were detected by an infrared beam in the recessed port.

### Electrophysiological recordings

For electrophysiological recordings, rats were tethered via a cable from their headstage to a commutator in the center of the chamber ceiling. Electrical signals and drinking events were collected using the OmniPlex system (Plexon). We recorded from the same location for two sessions if new neurons appeared on previously unrecorded channels. If multiple sessions for the same location were included in the analysis, the same channel was never included more than once. After the second recording in the same location, the drive was advanced 160 μm and recording resumed in the new location at minimum two days later to ensure settling of the tissue around the wires.

### Two bottle choice with optogenetic manipulation

For the optogenetic experiments, rats were habituated to being tethered to 200 μm core diameter patch cords (Doric Instruments) connected to a commutator (Doric Instruments) in turn connected to a 473 nm DPSS laser (Opto-Engine LLC). During testing rats were placed in a modified home cage that allowed the presentation of two individual bottles via ports on one wall of the homecage with the bottles hanging outside the cage. In daily 30-minute sessions, rats were presented with two possible solutions and allowed to freely drink. Licks made on each bottle were recorded using a custom-built lickometer system using Arduino and a capacitive MPR121 sensor (Adafruit Industries). The Arduino recorded licks in real time from each bottle, and one bottle each day was set as the active bottle such that the first lick made to that bottle each second would trigger a TTL pulse to a Master9 Stimulus Controller (AMPI) that dictated the duration and parameters of laser stimulation. For optoexcitation experiments, light was delivered for 1s at 20 Hz (5 ms ON, 50 ms OFF) and light output was calibrated to 8-12 mW from the end of the patchcord. For optoinhibition experiments, light was delivered continuously from the start of a lick bout until no lick was detected for 1s and light output was calibrated to 15-20 mW from the end of the patchcord. The order of testing, the side of the active bottle and the identity of solutions was counterbalanced. The weight of each bottle was recorded before and after each session and the rats were weighed before each session to identify the amount of solution consumed.

### Optogenetic intracranial self stimulation

Intracranial self-stimulation was conducted both in the two-bottle choice apparatus described above and in a standard MedAssociates operant chamber. For the two-bottle choice ICSS tests, rats were presented with either empty bottles or bottles containing water for a 30-minute session. One bottle, side counterbalanced across tests and rats, was designated as active such that the first lick on that bottle each second triggered a 1s, 20 Hz (5 ms ON, 50 ms OFF) train of 473 nm light bilaterally into the central amygdala with light output set at 8-12 mW from the end of the patchcord. Responses were recorded on the active and inactive bottle as well as the number of stimulations earned. For nosepoke ICSS, rats were placed into a MedAssociates operant chamber, connected to 473 nm lasers via patchcords and commutators and in a 1-hour session allowed to nosepoke in either of two ports. One port was designated as active where the first poke in that port each second delivered a 1 s, 20 Hz (5 ms ON, 50 ms OFF) train of 473 nm light bilaterally into the central amygdala with light output set at 8-12 mW from the end of the patchcord. Pokes into each port and stimulations earned were recorded by MedAssociates software.

### Electrophysiology data analysis

Isolation of individual units was performed using Offline Sorter (Plexon) by first manually selecting units based on clustering of waveforms. Units were then separated and refined using interspike interval distribution, cross-correlograms, and autocorrelograms. Any units that were not detectable for the entire session were not included in the study. Sorted units were exported to NeuroExplorer 3.0 (Nex Technologies) and MATLAB (Mathworks) for all subsequent analysis. Neurons were determined to be modulated by an event if the spike rate in a custom window (−0.5 to 0.5 s for port entries and port exits and 0 to 0.03 s for lick) following each event significantly differed from a 10 s baseline period according to a Wilcoxon signed-rank test (p < 0.05, two-tailed). Peri-stimulus time histograms (PSTHs) were constructed around event-related responses using 0.01 ms bins. The spiking activity of each neuron across these bins of the PSTH was smoothed using a half-normal filter (σ = 6.6) that used activity in previous, but not upcoming, bins. To visualize the normalized activity of neurons, the mean activity within each of the smoothed bins of the PSTH was transformed to a z-score as follows: (F_i_ – F_mean_)/ F_SD_, where F_i_ is the firing rate of the i^th^ bin of the PSTH, and F_mean_ and F_SD_ are the mean and SD of the firing rate of the 10 s baseline period. Color-coded maps and average traces of individual neurons’ activity were constructed based on these z-scores.

### Lick-modulation analysis

This analysis was restricted to licks emitted in bouts, i.e. with inter-lick intervals < 210ms. Distributions of spike phases (in radians) were computed for each neuron (neurons with less than 50 spikes in lick cycles were excluded) and non-uniformity was tested with Rayleigh test. Neurons with p<0.01 were considered lick-modulated. V test was used to test for non-uniformity of preferred firing phase distribution with a mean direction of 90 degrees (**Figure 2G** and **Supplement 2F**). The distributions of the preferred firing phases of lick-modulated neurons from ethanol and sucrose consuming rats were compared using the Kuiper test (circular analogue of the Kolmogorov-Smirnov test; **Supplement 2G**). Proportions of lick-modulated neurons in ethanol and sucrose consuming rats were compared using a z binomial proportion test (**Supplement 2H**).

### Statistical analysis

Data are presented as mean ± s.e.m. unless otherwise indicated in the text. Statistical analyses were performed using either MATLAB (Mathworks) or in Prism 8 (GraphPad). For electrophysiological data, statistical tests were performed on unsmoothed data. The specific tests performed are noted throughout the text and figure legends. For electrophysiological data we did not test for normality, but made use of nonparametric tests (two-sided Wilcoxon’s rank-sum and signed-rank tests). For optogenetic data we made use of two-way repeated measures ANOVA and post hoc tests were performed with Sidak’s method when appropriate and t-tests performed with Welch’s correction for unequal standard deviations between groups. For the inhibiton experiment we had an a priori hypothesis to conduct post hoc comparisons within each virus group across the two bottles based on our findings with optogenetic stimulation. Each optogenetic test was conducted only once per rat. Three eNpHR rats did not drink alcohol in any of the tests despite repeated efforts so their data was excluded in these cases, they still performed in all other experiments and were included in those tests.

## ACKNOWLEDGEMENTS

The authors have no biomedical financial interests or potential conflicts of interest. This research was supported by grants from the National Institutes of Health to K.M.F (F31 DA043136) and P.H.J (R01 AA026306 & R01 AA027213) and the National Science Foundation to D.J.O (DGE-1746891).

## SUPPLEMENTAL FIGURES

**FIGURE S1:**
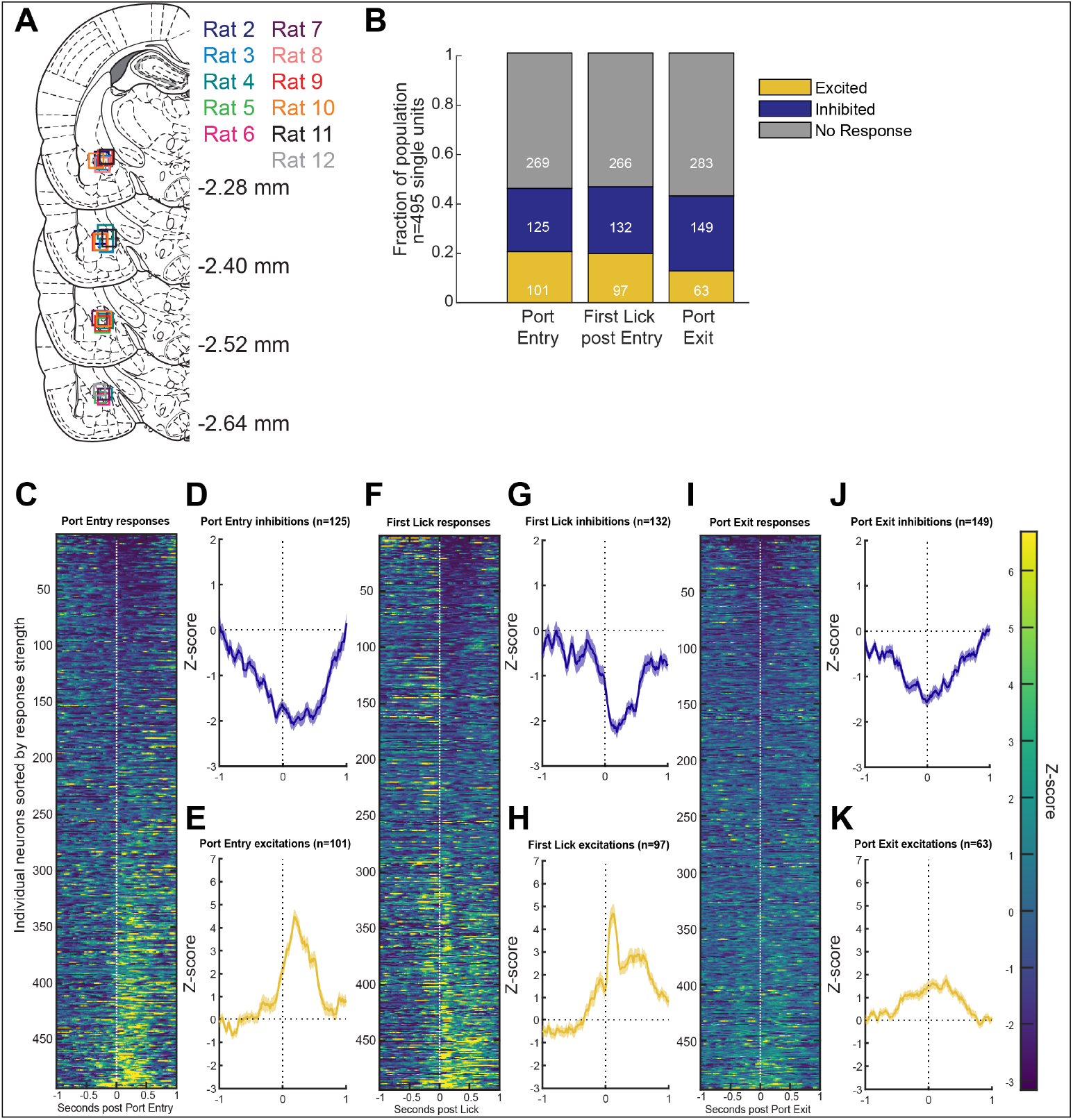
Identifying correlates of sucrose consumption in the central nucleus of the amygdala. **A)** Recreation of recording sites from each of the five rats. The task was identical to that in **Figure 1**. Rats made on average 50.91 ± 6.637 port entries and 1486 ± 212.4 licks **B)** Proportion of neurons significantly excited or inhibited by task-relevant events. There were no differences in the proportion of neurons excited or inhibited by port entries (X^2^ = 2.549, p = 0.1104), but more neurons were inhibited than excited by port exits (X^2^ = 34.887, p = 0.0001) and the first lick after a port entry (X^2^ = 5.349, p = 0.0207). In addition, more neurons were inhibited by the the first lick than port exit (X^2^ = 8.7837, p = 0.0124). **C)** Heatmap of z-scored responses for each neuron recorded sorted by the strength of excitation to port entry. **D)** Average z-scored response of all neurons that were identified as being significantly inhibited around port entry. **E)** Average z-scored response of all neurons that were identified as significantly excited around port entry. **F-H)** Same as **C-E** but for the first lick post port entry. **I-K)** Same as **C-E** but for port exit. Traces indicate mean z-scored response with overlaid bands indicating ± 1 standard error of the mean.

**FIGURE S2:**
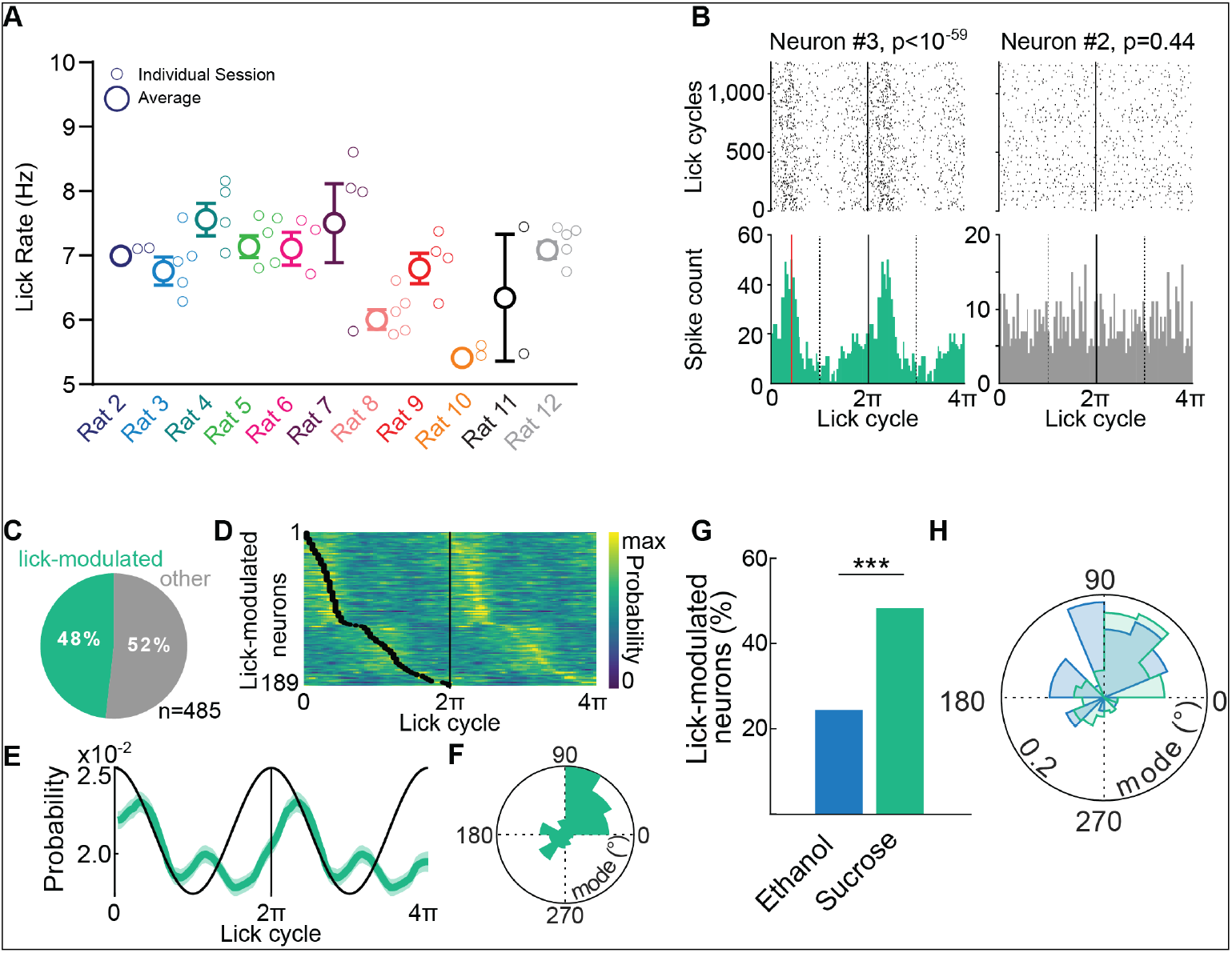
Central amygdala neurons are modulated by licks during the consumption of sucrose. **A)** Average lick rates during the consumption of 14.2% sucrose for each rat during recording sessions. Smaller symbols indicate lick rate for each individual session. **B)** Spike rasters (top) and histograms (bottom) during lick cycles of two example neurons recorded in the same session. A lick cycle is defined as the time between two consecutive contacts with the fluid delivery port (see Methods). The p-value of Rayleigh test is indicated. **C)** Proportion of neurons significantly modulated by licks (Rayleigh’s test with p-value < 0.01). **D)** Heat map of spike probability during lick cycles of lick-modulated neurons. Black dots indicate the preferred firing phases (i.e. modes). **E)** Average spike probability of lick-modulated neurons across lick cycles (mean±s.e.m.). **F)** Circular histogram of the preferred firing phases (V test against 90°, n=189 lick-modulated neurons, V_189_=53.43, p<10^−7^). G) The proportion of lick-modulated neurons is higher during sucrose consumption compared to ethanol (z binomial proportion test, p<10^−6^). **H)** The distributions of the preferred firing phases of lick-modulated neurons from ethanol and sucrose consuming rats are not significantly different (Kuiper test, k=2.1090.103, K=1.9931.103, p=0.1).

**FIGURE S3:**
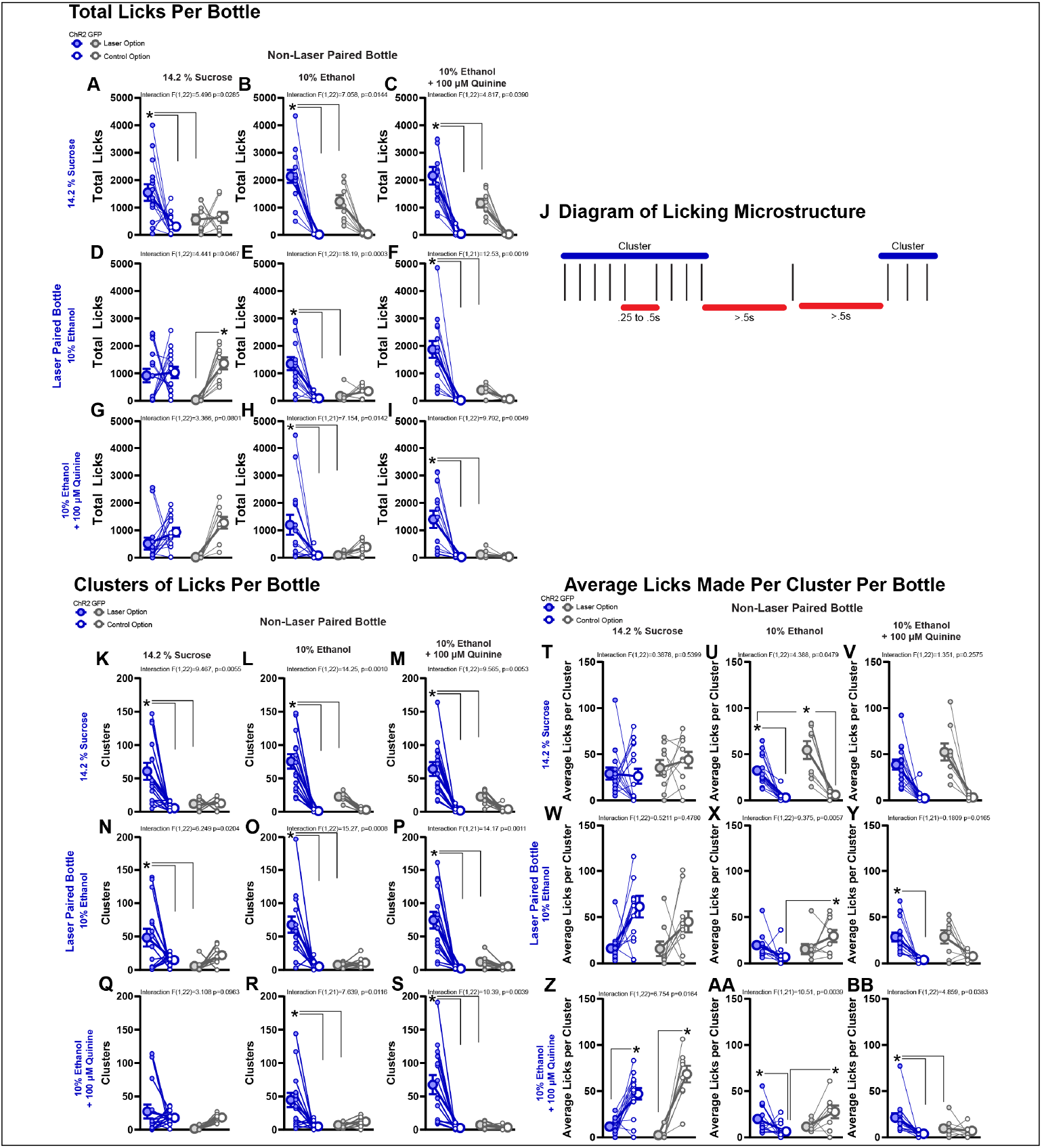
Microstructural analysis of consumption indicates optogenetic stimulation of the central amygdala enhances motivation to consume but not palatability of the laser-paired option. **A-I)** Total number of licks made on each bottle for each of the tests presented in **Figure 3**. Graphs are organized with the most valued option at the top and leftmost position and the least valued option at the bottom and rightmost position. Comparisons between bottles containing the same offer tile the diagonal, bottles above diagonal are tests in which the more valued option was blue-light paired and tests below the diagonal are when blue-light was paired with the less valued option **J)** Diagram of the licking microstructure used to separate out clusters. Clusters had at least three licks and a interlick interval of at least 500 ms. **K-S)** Same as **A-I** but the total number of clusters of licks made on each bottle. **T-BB)** Same as **A-I** but the average number of licks per cluster made on each bottle. Filled symbols indicate the bottle that resulted in blue light delivery, open symbols the other bottle that did not trigger any light delivery. Large symbols indicate group means ± 1 standard error of the mean and small symbols represent individual rats. * p<0.05 for post hoc comparisons made only when an interaction between virus and bottle was observed.

**FIGURE S4:**
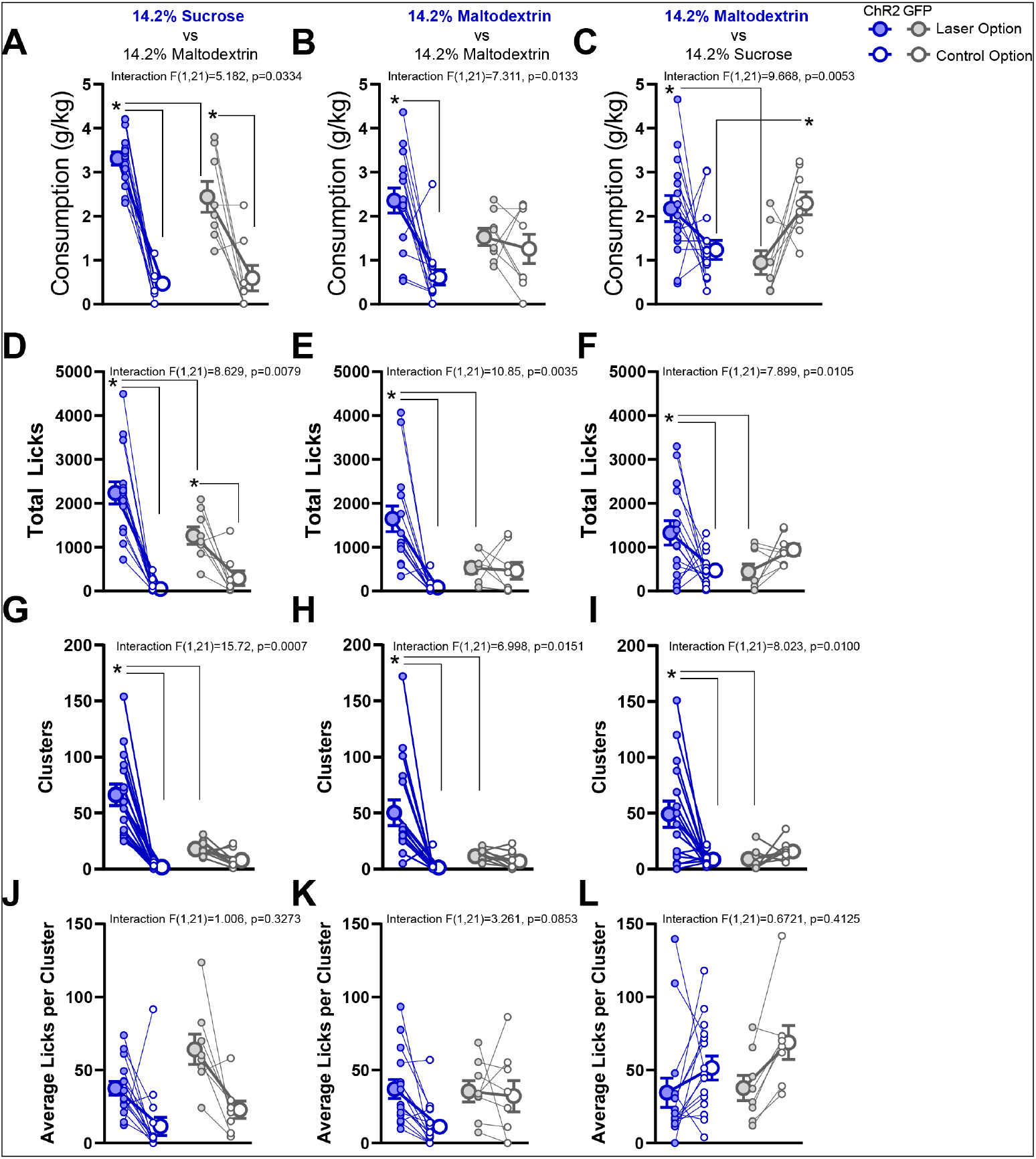
Optogenetic stimulation of the central amygdala during consumption can reverse preference between two isocaloric and objectively equal rewards. **A)** Consumption in g/kg in tests when sucrose consumption was laser-paired and maltodextrin was not. **B)** Consumption in g/kg in tests where one bottle containing maltodextrin was laser-paired and the other bottle with maltodextrin was not. **C)** Consumption in g/kg in tests when maltodextrin consumption was laser-paired and sucrose was not. **D-F)** Same as **A-C** but the number of licks made on each bottle. **G-I)** Same as **A-C** but the number of clusters of licks made on each bottle. **J-L)** Same as **A-C** but the average number of licks made per cluster for stimulation-paired option versus the non-paired option. Filled symbols indicate the bottle that resulted in blue light delivery, open symbols the other bottle that did not trigger any light delivery. Large symbols indicate group means ± 1 standard error of the mean and small symbols represent individual rats. * p<0.05 for post hoc comparisons made only when an interaction between virus and bottle was observed.

**FIGURE S5:**
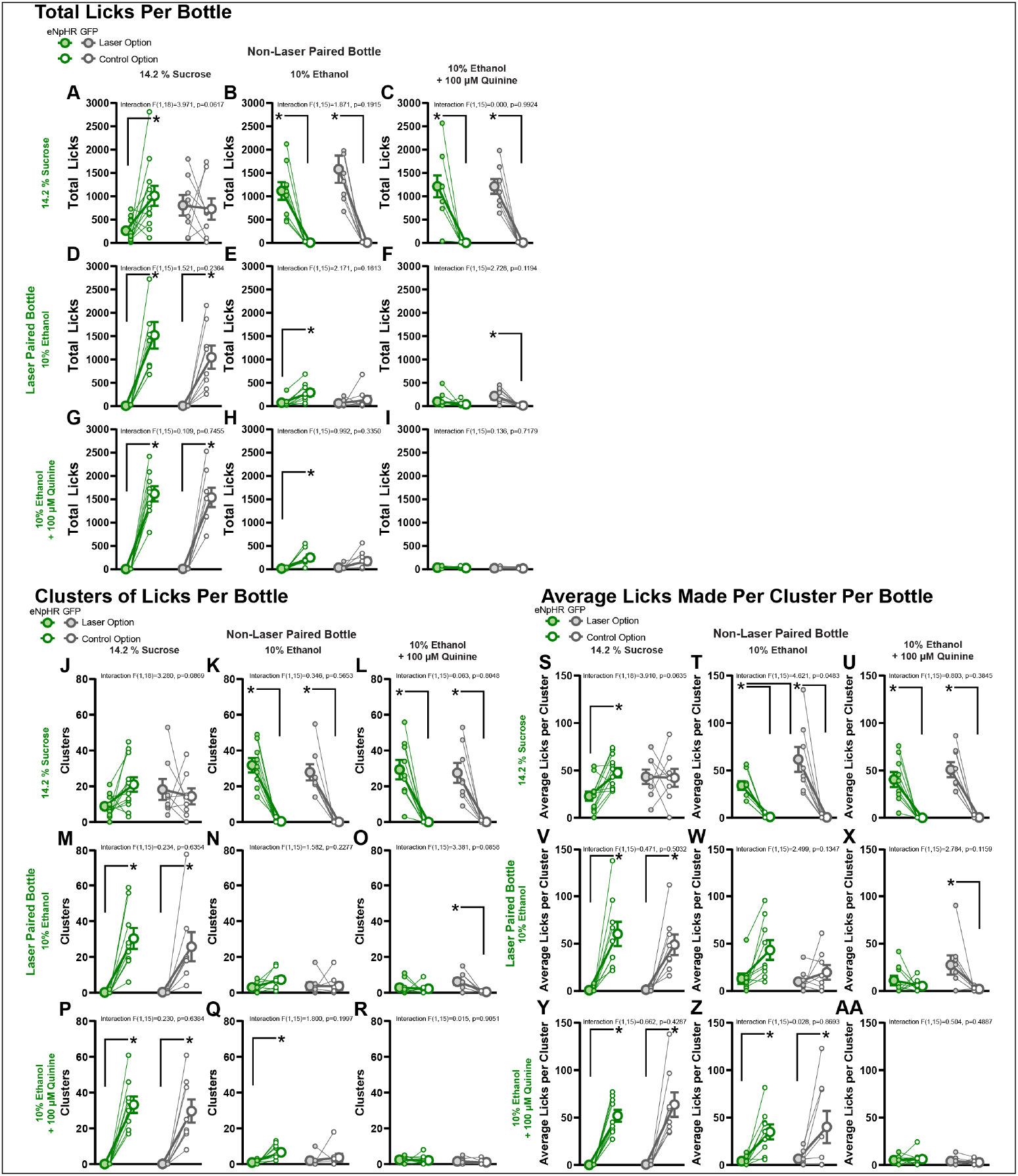
Microstructural analysis of consumption indicates optogenetic inhibition of the central amygdala does not suppress motivation to consume but suppresses the palatability of the laser-paired option. **A-I)** Total number of licks made on each bottle for each of the tests presented in **Figure 5.** Graphs are organized with the most valued option at the top and leftmost position and the least valued option at the bottom and rightmost position. Comparisons between bottles containing the same offer tile the diagonal, bottles above diagonal are tests in which the more valued option was green-light paired and tests below the diagonal are when green-light was paired with the less valued option **J-R)** Same as **A-I** but the total number of clusters of licks made on each bottle. **S-AA)** Same as **A-I** but the average number of licks per cluster made on each bottle. Filled symbols indicate the bottle that resulted in green light delivery, open symbols the other bottle that did not trigger any light delivery. Large symbols indicate group means ± 1 standard error of the mean and small symbols represent individual rats. * p<0.05 for post hoc comparisons.

**FIGURE S6:**
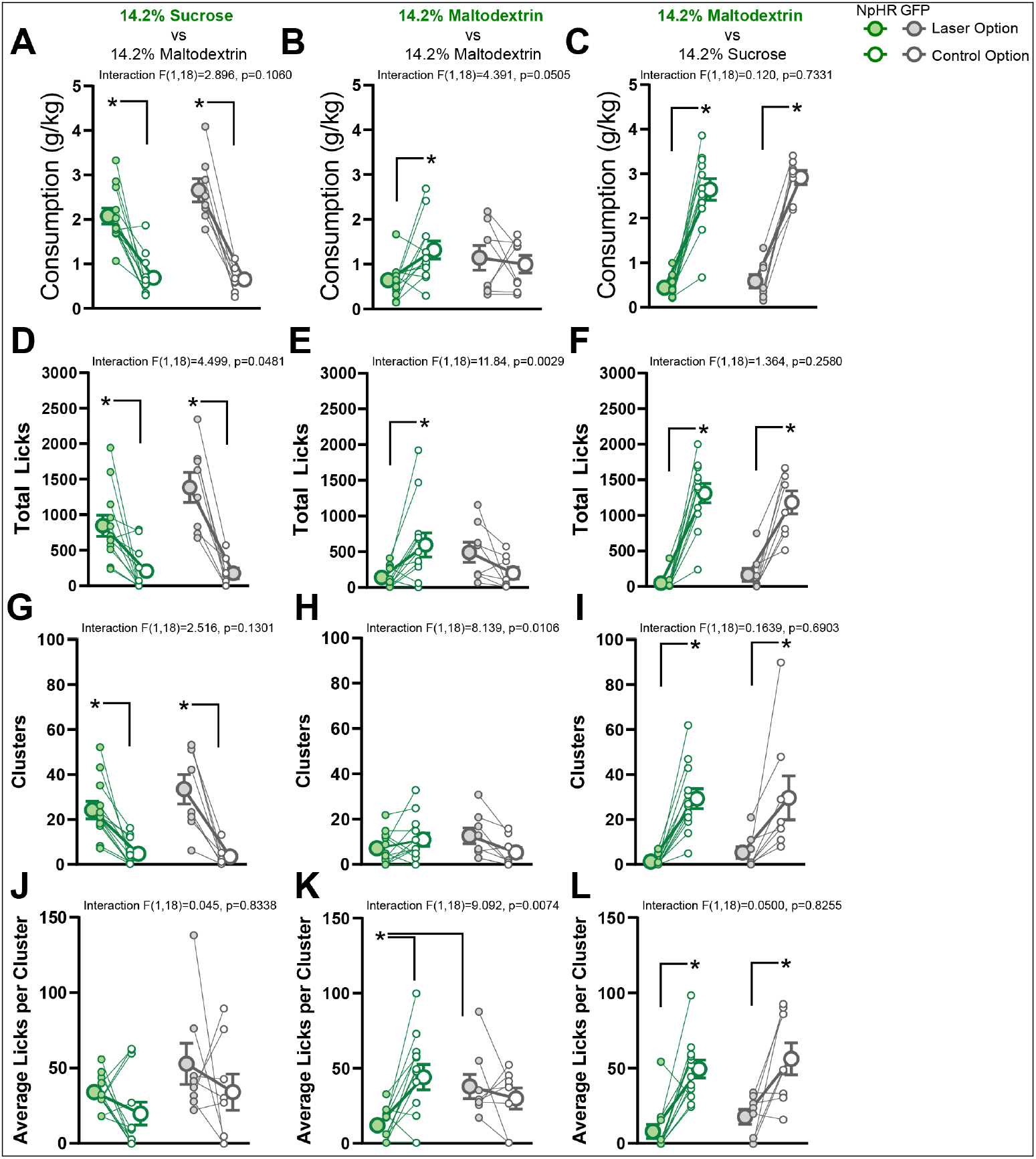
Optogenetic inhibition of the central amygdala during consumption cannot reverse preference between two isocaloric and palatable rewards. **A)** Consumption in g/kg in tests when sucrose consumption was laser-paired and maltodextrin was not. **B)** Consumption in g/kg in tests where one bottle containing maltodextrin was laser-paired and the other bottle with maltodextrin was not. **C)** Consumption in g/kg in tests when maltodextrin consumption was laser-paired and sucrose was not. **D-F)** Same as **A-C** but the number of licks made on each bottle. **G-I)** Same as **A-C** but the number of clusters of licks made on each bottle. **J-L)** Same as **A-C** but the average number of licks made per cluster for inhibition-paired option versus the non-paired option. Filled symbols indicate the bottle that resulted in green light delivery, open symbols the other bottle that did not trigger any light delivery. Large symbols indicate group means ± 1 standard error of the mean and small symbols represent individual rats. * p<0.05 for post hoc comparisons.

## REFERENCES

1. Koob, G. F. & Volkow, N. D. Neurobiology of addiction: a neurocircuitry analysis. Lancet Psychiatry 3, 760–773 (2016).

2. Robinson, T. E. & Berridge, K. C. The neural basis of drug craving: an incentive-sensitization theory of addiction. Brain Research. Brain Research Reviews 18, 247–291 (1993).

3. Sweis, B. M., Thomas, M. J. & Redish, A. D. Beyond simple tests of value: measuring addiction as a heterogeneous disease of computation-specific valuation processes. Learn. Mem. 25, 501–512 (2018).

4. Augier, E. et al. A molecular mechanism for choosing alcohol over an alternative reward. Science 360, 1321–1326 (2018).

5. Foster, K. L. et al. GABA(A) and opioid receptors of the central nucleus of the amygdala selectively regulate ethanol-maintained behaviors. Neuropsychopharmacology 29, 269–284 (2004).

6. Mahler, S. V. & Berridge, K. C. Which cue to ‘want?’ Central amygdala opioid activation enhances and focuses incentive salience on a prepotent reward cue. The Journal of Neuroscience 29, 6500–6513 (2009).

7. Pelloux, Y., Minier-Toribio, A., Hoots, J. K., Bossert, J. M. & Shaham, Y. Opposite Effects of Basolateral Amygdala Inactivation on Context-Induced Relapse to Cocaine Seeking after Extinction versus Punishment. J. Neurosci. 38, 51–59 (2018).

8. Roberts, A. J., Cole, M. & Koob, G. F. Intra-amygdala muscimol decreases operant ethanol self-administration in dependent rats. Alcohol. Clin. Exp. Res. 20, 1289–1298 (1996).

9. Robinson, M. J. F., Warlow, S. M. & Berridge, K. C. Optogenetic excitation of central amygdala amplifies and narrows incentive motivation to pursue one reward above another. The Journal of Neuroscience 34, 16567–16580 (2014).

10. Tom, R. L., Ahuja, A., Maniates, H., Freeland, C. M. & Robinson, M. J. F. Optogenetic activation of the central amygdala generates addiction-like preference for reward. Eur. J. Neurosci. 50, 2086– 2100 (2019).

11. Torruella-Suárez, M. L. et al. Manipulations of Central Amygdala Neurotensin Neurons Alter the Consumption of Ethanol and Sweet Fluids in Mice. J. Neurosci. 40, 632–647 (2020).

12. Warlow, S. M., Naffziger, E. E. & Berridge, K. C. The central amygdala recruits mesocorticolimbic circuitry for pursuit of reward or pain. Nat Commun 11, 2716 (2020).

13. Warlow, S. M., Robinson, M. J. F. & Berridge, K. C. Optogenetic Central Amygdala Stimulation Intensifies and Narrows Motivation for Cocaine. J. Neurosci. 37, 8330–8348 (2017).

14. Baumgartner, H. M., Granillo, M., Schulkin, J. & Berridge, K. C. Corticotropin releasing factor (CRF) systems: Promoting cocaine pursuit without distress via incentive motivation. PLoS One 17, e0267345 (2022).

15. Vandaele, Y. & Ahmed, S. H. Habit, choice, and addiction. Neuropsychopharmacology 46, 689–698 (2021).

16. Ahmed, S. H., Lenoir, M. & Guillem, K. Neurobiology of addiction versus drug use driven by lack of choice. Current Opinion in Neurobiology 23, 581–587 (2013).

17. Bienkowski, M. S. & Rinaman, L. Common and distinct neural inputs to the medial central nucleus of the amygdala and anterior ventrolateral bed nucleus of stria terminalis in rats. Brain Struct Funct 218, 187–208 (2013).

18. Norgren, R. Taste pathways to hypothalamus and amygdala. J. Comp. Neurol. 166, 17–30 (1976).

19. Ottersen, O. P. & Ben-Ari, Y. Afferent connections to the amygdaloid complex of the rat and cat. I. Projections from the thalamus. J. Comp. Neurol. 187, 401–424 (1979).

20. Yasoshima, Y., Shimura, T. & Yamamoto, T. Single unit responses of the amygdala after conditioned taste aversion in conscious rats. Neuroreport 6, 2424–2428 (1995).

21. Douglass, A. M. et al. Central amygdala circuits modulate food consumption through a positive-valence mechanism. Nat. Neurosci. 20, 1384–1394 (2017).

22. Hardaway, J. A. et al. Central Amygdala Prepronociceptin-Expressing Neurons Mediate Palatable Food Consumption and Reward. Neuron 102, 1037–1052.e7 (2019).

23. Shabel, S. J. & Janak, P. H. Substantial similarity in amygdala neuronal activity during conditioned appetitive and aversive emotional arousal. Proceedings of the National Academy of Sciences of the United States of America 106, 15031–15036 (2009).

24. Nishijo, H., Uwano, T., Tamura, R. & Ono, T. Gustatory and multimodal neuronal responses in the amygdala during licking and discrimination of sensory stimuli in awake rats. J Neurophysiol 79, 21–36 (1998).

25. Yang, T. et al. Plastic and stimulus-specific coding of salient events in the central amygdala. Nature 616, 510–519 (2023).

26. Paton, J. J., Belova, M. A., Morrison, S. E. & Salzman, C. D. The primate amygdala represents the positive and negative value of visual stimuli during learning. Nature 439, 865–870 (2006).

27. Wang, L. et al. The coding of valence and identity in the mammalian taste system. Nature 558, 127–131 (2018).

28. Kim, J., Zhang, X., Muralidhar, S., LeBlanc, S. A. & Tonegawa, S. Basolateral to central amygdala neural circuits for appetitive behaviors. Neuron 93, 1464–1479.e5 (2017).

29. Gilpin, N. W. & Roberto, M. Neuropeptide modulation of central amygdala neuroplasticity is a key mediator of alcohol dependence. Neurosci Biobehav Rev 36, 873–888 (2012).

30. Gilpin, N. W., Herman, M. A. & Roberto, M. The central amygdala as an integrative hub for anxiety and alcohol use disorders. Biological Psychiatry 77, 859–869 (2015).

31. Roberto, M., Kirson, D. & Khom, S. The Role of the Central Amygdala in Alcohol Dependence. Cold Spring Harb Perspect Med 11, a039339 (2021).

32. Ron, D. & Barak, S. Molecular mechanisms underlying alcohol-drinking behaviours. Nat. Rev. Neurosci. 17, 576–591 (2016).

33. Egervari, G., Siciliano, C. A., Whiteley, E. L. & Ron, D. Alcohol and the brain: from genes to circuits. Trends in Neurosciences 44, 1004–1015 (2021).

34. Barak, S. et al. Disruption of alcohol-related memories by mTORC1 inhibition prevents relapse. Nat. Neurosci. 16, 1111–1117 (2013).

35. Carnicella, S., Ron, D. & Barak, S. Intermittent ethanol access schedule in rats as a preclinical model of alcohol abuse. Alcohol (Fayetteville, N.Y.) 48, 243–252 (2014).

36. Wise, R. A. Voluntary ethanol intake in rats following exposure to ethanol on various schedules. Psychopharmacologia 29, 203–210 (1973).

37. Naneix, F., Peters, K. Z. & McCutcheon, J. E. Investigating the Effect of Physiological Need States on Palatability and Motivation Using Microstructural Analysis of Licking. Neuroscience (2019) doi:10.1016/j.neuroscience.2019.10.036.

38. Ottenheimer, D., Richard, J. M. & Janak, P. H. Ventral pallidum encodes relative reward value earlier and more robustly than nucleus accumbens. Nat Commun 9, 4350 (2018).

39. Gutierrez, R., Simon, S. A. & Nicolelis, M. A. L. Licking-induced synchrony in the taste-reward circuit improves cue discrimination during learning. J. Neurosci. 30, 287–303 (2010).

40. Hopf, F. W. & Lesscher, H. M. B. Rodent models for compulsive alcohol intake. Alcohol 48, 253–264 (2014).

41. Spanagel, R., Hölter, S. M., Allingham, K., Landgraf, R. & Zieglgänsberger, W. Acamprosate and alcohol: I. Effects on alcohol intake following alcohol deprivation in the rat. European Journal of Pharmacology 305, 39–44 (1996).

42. Wolffgramm, J. & Heyne, A. From controlled drug intake to loss of control: the irreversible development of drug addiction in the rat. Behav. Brain Res. 70, 77–94 (1995).

43. Spector, A. C. & St John, S. J. Role of taste in the microstructure of quinine ingestion by rats. Am. J. Physiol. 274, R1687-1703 (1998).

44. Spector, A. C., Klumpp, P. A. & Kaplan, J. M. Analytical issues in the evaluation of food deprivation and sucrose concentration effects on the microstructure of licking behavior in the rat. Behavioral Neuroscience 112, 678–694 (1998).

45. Davis, J. D. & Perez, M. C. Food deprivation- and palatability-induced microstructural changes in ingestive behavior. Am. J. Physiol. 264, R97-103 (1993).

46. Keiflin, R., Reese, R. M., Woods, C. A. & Janak, P. H. The orbitofrontal cortex as part of a hierarchical neural system mediating choice between two good options. The Journal of Neuroscience 33, 15989–15998 (2013).

47. Nissenbaum, J. W. & Sclafani, A. Qualitative differences in polysaccharide and sugar tastes in the rat: A two-carbohydrate taste model. Neuroscience & Biobehavioral Reviews 11, 187–196 (1987).

48. Ottenheimer, D. J. et al. A quantitative reward prediction error signal in the ventral pallidum. Nat Neurosci 23, 1267–1276 (2020).

49. Servonnet, A., Hernandez, G., El Hage, C., Rompré, P.-P. & Samaha, A.-N. Optogenetic Activation of the Basolateral Amygdala Promotes Both Appetitive Conditioning and the Instrumental Pursuit of Reward Cues. J. Neurosci. 40, 1732– 1743 (2020).

50. Steinberg, E. E. et al. Amygdala-Midbrain Connections Modulate Appetitive and Aversive Learning. Neuron 106, 1026–1043.e9 (2020).

51. Davis, M., Walker, D. L., Miles, L. & Grillon, C. Phasic vs sustained fear in rats and humans: role of the extended amygdala in fear vs anxiety. Neuropsychopharmacology 35, 105–135 (2010).

52. Carter, M. E., Han, S. & Palmiter, R. D. Parabrachial calcitonin gene-related peptide neurons mediate conditioned taste aversion. J. Neurosci. 35, 4582–4586 (2015).

53. Schiff, H. C. et al. An Insula-Central Amygdala Circuit for Guiding Tastant-Reinforced Choice Behavior. J. Neurosci. 38, 1418–1429 (2018).

54. Bloodgood, D. W. et al. Kappa opioid receptor and dynorphin signaling in the central amygdala regulates alcohol intake. Mol. Psychiatry (2020) doi:10.1038/s41380-020-0690-z.

55. Cai, H., Haubensak, W., Anthony, T. E. & Anderson, D. J. Central amygdala PKC-δ(+) neurons mediate the influence of multiple anorexigenic signals. Nat. Neurosci. 17, 1240–1248 (2014).

56. Fadok, J. P., Markovic, M., Tovote, P. & Lüthi, A. New perspectives on central amygdala function. Curr. Opin. Neurobiol. 49, 141–147 (2018).

57. Han, W. et al. Integrated control of predatory hunting by the central nucleus of the amygdala. Cell 168, 311–324.e18 (2017).

58. Galaverna, O. G. et al. Lesions of the central nucleus of the amygdala I: Effects on taste reactivity, taste aversion learning and sodium appetite. Behavioural Brain Research 59, 11–17 (1993).

59. Han, W. et al. Integrated Control of Predatory Hunting by the Central Nucleus of the Amygdala. Cell 168, 311–324.e18 (2017).

60. Van Daele, D. J., Fazan, V. P. S., Agassandian, K. & Cassell, M. D. Amygdala connections with jaw, tongue and laryngo-pharyngeal premotor neurons. Neuroscience 177, 93–113 (2011).

61. de Guglielmo, G. et al. Recruitment of a Neuronal Ensemble in the Central Nucleus of the Amygdala Is Required for Alcohol Dependence. J. Neurosci. 36, 9446–9453 (2016).

62. de Guglielmo, G. et al. Inactivation of a CRF-dependent amygdalofugal pathway reverses addiction-like behaviors in alcohol-dependent rats. Nat Commun 10, 1238 (2019).

63. Koob, G. F. et al. Addiction as a stress surfeit disorder. Neuropharmacology 76 Pt B, 370–382 (2014).

64. Venniro, M. et al. Abstinence-dependent dissociable central amygdala microcircuits control drug craving. Proceedings of the National Academy of Sciences 117, 8126–8134 (2020).

65. Radke, A. K., Sneddon, E. A., Frasier, R. M. & Hopf, F. W. Recent Perspectives on Sex Differences in Compulsion-Like and Binge Alcohol Drinking. Int J Mol Sci 22, 3788 (2021).

